# Mutual information networks reveal evolutionary relationships within the influenza A virus polymerase

**DOI:** 10.1101/2023.02.16.528850

**Authors:** Sarah Arcos, Alvin X. Han, Aartjan J. W. te Velthuis, Colin A. Russell, Adam S. Lauring

## Abstract

The influenza A (IAV) RNA polymerase is an essential driver of IAV evolution. Mutations that the polymerase introduces into viral genome segments during replication are the ultimate source of genetic variation, including within the three subunits of the IAV polymerase (PB2, PB1, and PA). Evolutionary analysis of the IAV polymerase is complicated, because changes in mutation rate, replication speed, and drug resistance involve epistatic interactions among its subunits. In order to study the evolution of the human seasonal H3N2 polymerase since the 1968 pandemic, we identified pairwise evolutionary relationships among ∼7000 H3N2 polymerase sequences using mutual information (MI), which measures the information gained about the identity of one residue when a second residue is known. To account for uneven sampling of viral sequences over time, we developed a weighted MI metric (wMI) and demonstrate that wMI outperforms raw MI through simulations using a well-sampled SARS-CoV-2 dataset. We then constructed wMI networks of the H3N2 polymerase to extend the inherently pairwise wMI statistic to encompass relationships among larger groups of residues. We included HA in the wMI network to distinguish between functional wMI relationships within the polymerase and those potentially due to hitchhiking on antigenic changes in HA. The wMI networks reveal coevolutionary relationships among residues with roles in replication and encapsidation. Inclusion of HA highlighted polymerase-only subgraphs containing residues with roles in the enzymatic functions of the polymerase and host adaptability. This work provides insight into the factors that drive and constrain the rapid evolution of influenza viruses.

## Introduction

The evolution of influenza A viruses is constrained by epistatic interactions that limit viral exploration of sequence space (Lyons and Lauring 2018). Thus, epistasis can alter how influenza A viruses evade our two primary pharmaceutical interventions – vaccines and antiviral drugs. While most RNA viruses encode a single subunit polymerase, influenza A viruses (IAVs) express a heterotrimeric polymerase (Te Velthuis et al. 2021). This complex, consisting of polymerase basic protein 2 (PB2), polymerase basic protein 1 (PB1), and polymerase acidic protein (PA), works with nucleoprotein (NP) to bind viral RNA and carry out transcription and genome replication (Te Velthuis et al. 2021). Complex relationships between all three subunits determine the functions of the IAV polymerase. Furthermore, recent studies indicate that epistatic relationships within the IAV polymerase can manifest as a genetic barrier to drug resistance (Bloom et al. 2010; Pauly et al. 2017; Goldhill et al. 2018).

Epistasis, a non-additive fitness relationship between mutations, can occur due to structural and/or functional interactions. One indicator of protein epistasis is coevolution between residues, which can be measured when enough sequence data over evolutionary time is available. Inferring epistasis from coevolution assumes that the co-selection of two or more mutations arises as a result of a positive epistatic relationship between these mutations (Dunn et al. 2008). Existing approaches for measuring coevolution between protein residues tend to rely on phylogenetic inference (Yeang and Haussler 2007; Gong et al. 2013), which requires significant computational resources and is subject to issues with model mis-specification (e.g. different models can result in different trees and thus different estimates of coevolution) (Dutheil 2012).

In contrast, methods based on information theory do not require model fitting and can detect a broader range of relationships. For example, mutual information (MI) (Shannon 1948), which measures the amount of information shared between two random variables, has been used to identify co-evolving residues in proteins (Dunn et al. 2008; Dutheil 2012). Substantial effort has been spent in refining MI to predict protein structure by identifying residue contacts (Weigt et al. 2009; Morcos et al. 2011; Kamisetty et al. 2013; Figliuzzi et al. 2016). However, IAV polymerase evolution is likely driven by factors beyond structural contacts. For instance, protein allostery, RNA-protein interactions, RNA-RNA interactions, and interactions with cellular binding partners (including the ribosome and tRNAs) can all influence epistatic relationships within the IAV polymerase (Pflug et al. 2014; Dadonaite et al. 2019; Kim et al. 2020).

While information theory provides simple and interpretable tools for studying co-evolutionary relationships using sequencing data, there are several biases that need to be addressed prior to its application. First, these measures do not account for uneven sampling across categories or time. Second, they are limited to identifying pairwise interactions. Third, they do not address the possibility of genetic hitchhiking. Here we present solutions to these three problems and use the improved MI calculation to identify coevolutionary relationships within the H3N2 polymerase complex.

## Results

When applied to a multiple sequence alignment, MI quantifies the amount of information (measured as Shannon entropy) gained about one random variable (*H*(*a*), the entropy of site *a*) by observing a second random variable (*H*(*b*), the entropy of site *b*) (Shannon 1948) (Equations 1 and 2).

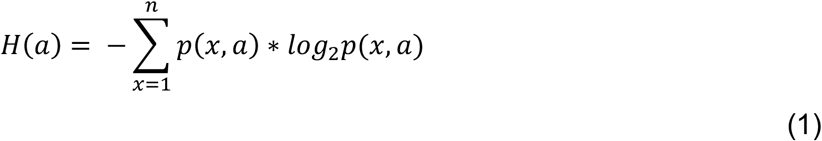

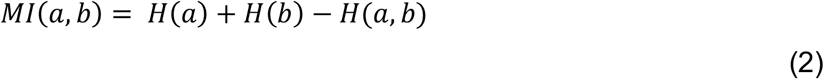

Where n is the number of columns in the alignment, *p*(*x, a*) is the frequency of a given amino acid, *x*, in site *a*, and *H*(*a, b*) is the joint entropy of *a* and *b* (calculated using di- residue frequencies).

Thus, MI quantifies how much easier it would be to predict the identity of an observed residue in one site if the identity of the residue in a second site is known. Importantly, MI is zero when the compared sites are completely conserved or completely randomly assorting.

### Weighted MI corrects for uneven sequence sampling over time

To quantify the MI between residues in the H3N2 polymerase, we first generated a joint multiple sequence alignment (MSA) of all complete H3N2 polymerase sequences (PB2, PB1, and PA) available on GISAID from 1968 to 2015. There were increasing numbers of IAV genomes available in recent years as sequencing technology advanced and surveillance infrastructure expanded; more H3N2 genomes were sequenced in 2015 than in the first five decades of H3N2 infections combined (Figure 1A). Because MI is calculated from the frequencies of a pair of random variables (Equations 1 and 2), calculations of entropy and MI will be more influenced by heavily sampled years. However, the skewed sampling over time will only alter these calculations if the MI (and entropy) change over time for residues in the IAV polymerase. We used a sliding-window approach to discover that the MI of H3N2 polymerase residues is not constant over time (Figure 1B). Therefore, calculations of MI across our entire dataset that do not account for the uneven sampling over time will be inflated for residues with high MI in recent years (e.g., PB2-590 and PB1-709, Figure 1B) and deflated for residues with high MI in earlier years (e.g., PA-350 and PB1-469, Figure 1B).

**Figure 1.**
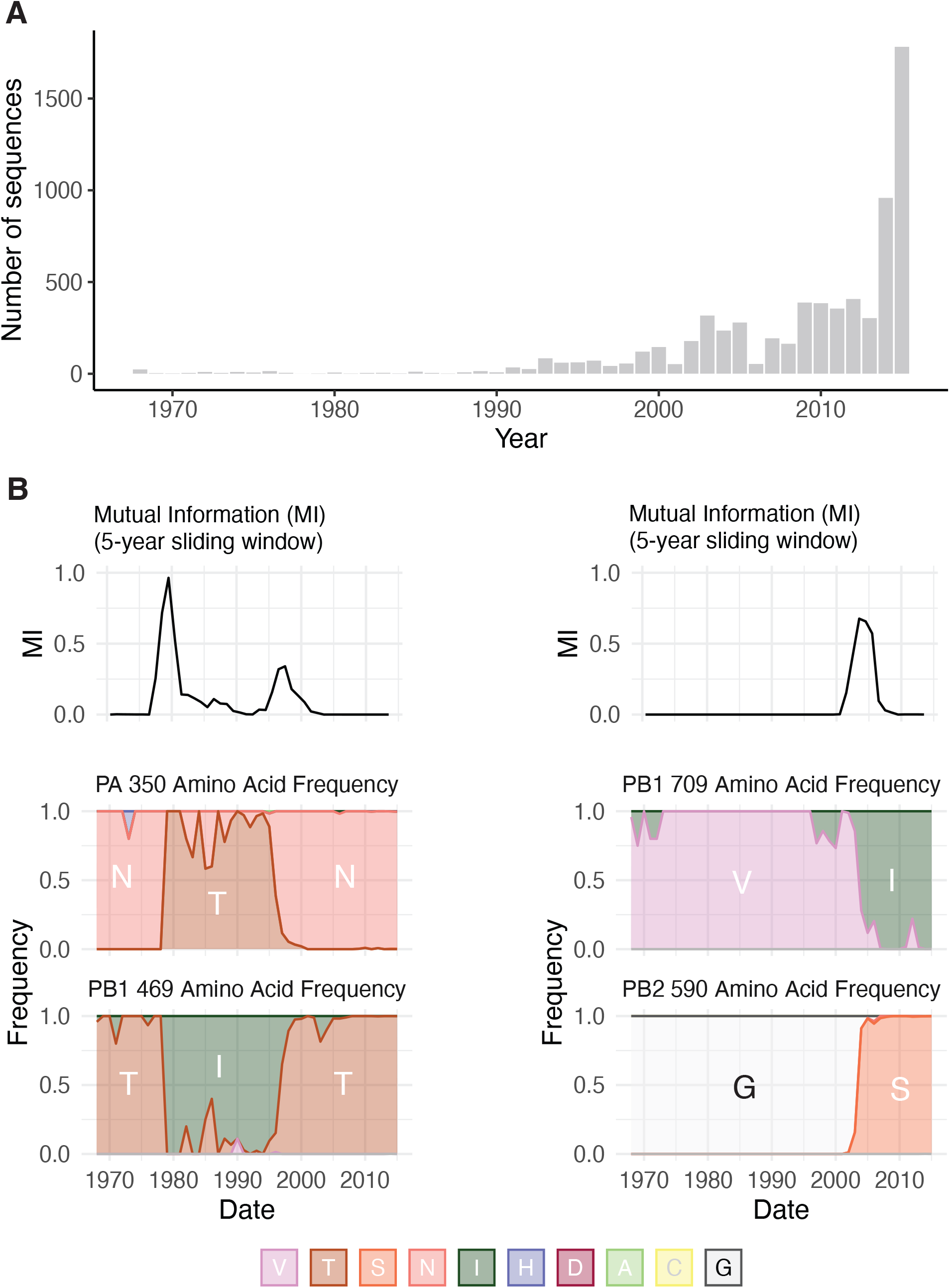
Uneven sampling of H3N2 polymerase sequences over time influences Shannon entropy and mutual information. (A) The distribution of complete H3N2 polymerase sequences on GISAID per year between 1968 and 2015. (B) Upper panels, Sliding window analysis of mutual information (MI) for residue pairs PA-350/PB1-469 and PB2-590/PB1-709. Sliding windows were constructed with a width of 5-years and a slide-length of 1 year. Lower panels, Plots of the frequency of amino acids for each residue over the period 1968 – 2015.

We accounted for the uneven sampling over time by creating weighted entropy and MI metrics. Previously, MI metrics have been developed that re-weight sequences in an MSA according to how many other sequences in the MSA exhibit similarity (e.g., Hamming distance) above a predefined threshold (Morcos et al. 2011). In our case, similarity re-weighting presents two issues. First, MI and sequence similarity are not independent and as such, re-weighting by one value will confound estimates of the other. Second, the distribution of similar sequences in our dataset contains essential information about selection and evolution that we want to capture in our calculation of MI. Thus, we designed new weighted entropy and MI metrics based on inverse probability weighting. Here, we used the weighted average of the residue frequencies (or di-residue frequencies) over each unit of time (e.g., year, month) to calculate the entropy and mutual information (Equation 3).

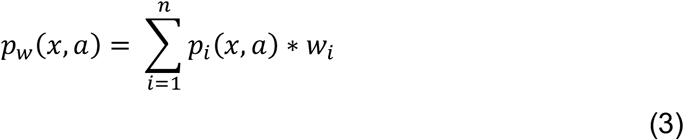

Where *n* is the number of time units and *w*_*i*_ is the weight for a given unit time.

We chose to apply the weighting procedure directly to the residue frequencies rather than the resulting entropy or MI to avoid overlooking years in which there is no residue variation (i.e., years where the entropy or MI are zero). We use “wMI” to refer to the weighted MI.

In an ideal scenario, the weight for each unit of time would be proportional to the number of virus infections per unit of time, as this would be best correlated with the amount of evolution. However, surveillance data from the early decades of H3N2 circulation is also variable and incomplete. Therefore, we evaluated how equal-weighting (Eq. 4) of each unit of time would compare to either weighting by disease incidence (Eq. 5) or no weighting using a dataset of SARS-CoV-2 spike RBD protein sequences generated by our laboratory in 2021 and 2022 (Valesano, Fitzsimmons, et al. 2021; Valesano, Rumfelt, et al. 2021).

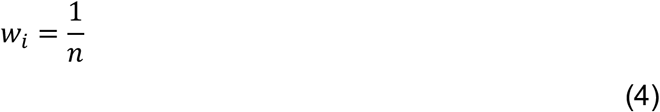

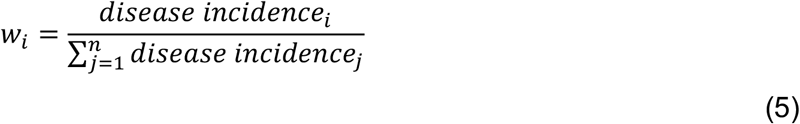

The original spike protein dataset is evenly sampled over each month with respect to disease incidence (Figure 2A) (https://www.michigan.gov/coronavirus/stats). We first generated 100 samples with replacement of the Spike MSA to simulate the uneven sampling present in the H3N2 polymerase MSA (Figure 2B, compare to Figure 1A) (see Methods). We then assessed the ability of wMI to correct for the simulated uneven sampling by calculating the unweighted, equal-weighted, and incidence-weighted wMIs for each sample and comparing these values to the MIs calculated from the original spike dataset. We found that incidence-weighted and equal-weighted wMIs closely approximated the MIs from the original spike dataset (incidence- weighted mean ρ = 0.985, 95% CI: 0.964 – 0.995; equal-weighted mean ρ = 0.971, 95% CI: 0.956 – 0.980) (Figure 2C). Moreover, both weighting procedures significantly outperformed the unweighted MI (mean ρ = 0.904, 95% CI: 0.841 – 0.945). This analysis shows that wMI calculated with equal-weighting or incidence-weighting yields improved calculations of the true MI for datasets that are unevenly sampled over time. Because we do not have good incidence data for H3N2 infections over time, we used equal weighting to calculate the pairwise wMI scores within the H3N2 polymerase.

**Figure 2.**
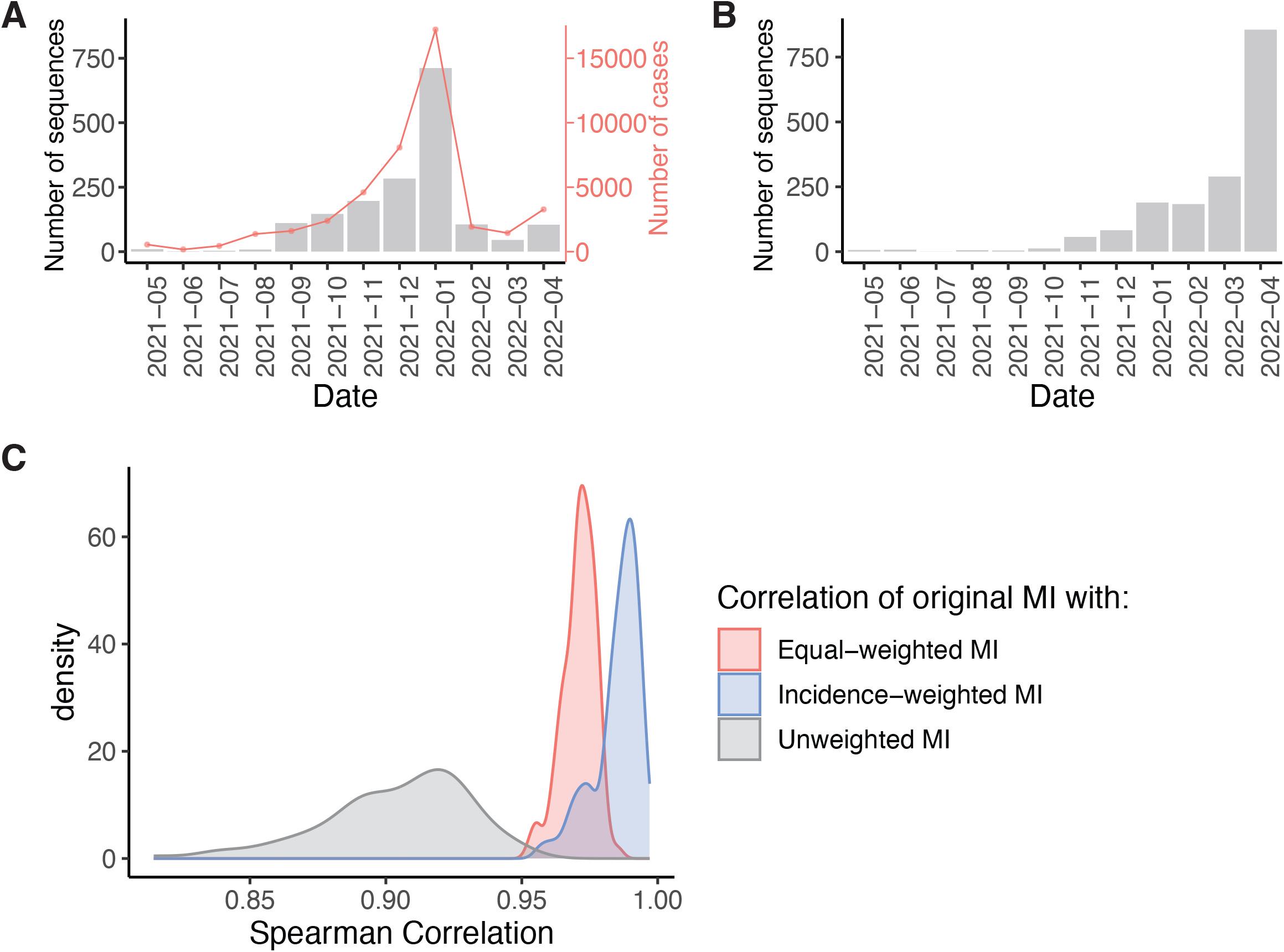
Re-weighting of amino acid frequencies improves MI estimates for unevenly sampled data. (A) The distribution of SARS-CoV-2 Spike RBD sequences generated by our laboratory per month between May 1st, 2021, and April 30th, 2021 from Washtenaw County, MI. The red line shows the number of confirmed COVID-19 cases in Washtenaw County, MI over the same time period. (B) Representative distribution of sampled Spike RBD sequences used to simulate the uneven sampling of H3N2 polymerase sequences (see Figure 1A). (C) The distribution of Spearman correlation coefficients between the MI from the original Spike dataset and the unweighted, equal-weighted, or incidence-weighted MI of 100 sampled datasets.

### Correcting wMI for the influence of phylogenetic relationships

Entropy and MI assume that all observations in a dataset are independent (Shannon 1948). However, as essentially every H3N2 polymerase sequence (since the reassortment event in 1968 that introduced avian PB1) has shared ancestry, this assumption is strongly violated (Dutheil 2012). The average-product correction (APC) devised by Gloor et al. corrects for phylogenetic relationships by estimating the background MI signal due to non-independence (Dunn et al. 2008). This is accomplished by calculating the mean MI for each member of a residue pair and for the dataset as a whole (Equation 6), which therefore assumes that the true number of coevolving amino acid pairs is a tiny fraction of the total possible pairs in the MSA.

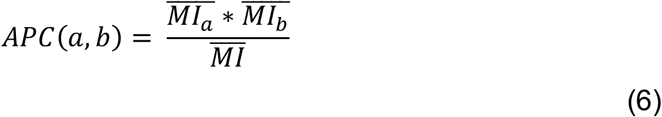

The corrected MI (or corrected wMI) for a given pair is calculated by subtracting the APC.

### wMI reveals coevolutionary relationships among mutations crucial for host range expansion

We next investigated pairwise coevolutionary relationships within the H3N2 polymerase complex. Calculating an equal-weighted wMI by influenza season would only be possible for one fifth of the time period covered by the H3N2 polymerase dataset, as we only have reliable collection month information for sequences after ∼2003. Therefore, we chose to weight across collection year (rather than season, month, or week), because that is the highest level of precision across all sequence metadata in our dataset.

We first investigated whether the top wMI scores capture known relationships within the H3N2 polymerase. For example, the PB2 627 residue is known to mediate adaptation to mammalian hosts, and mutations in or near this residue often occur during host range expansion to restore ANP32A binding and improve viral replication (Subbarao et al. 1993). Among the top wMI pairs (z-score > 4), 53 residues coevolve with PB2-627. We identified the top five residues paired with PB2-627 by wMI: PB2-44, PB2-199, PB2-591, PB2-645, and PA-268, and then plotted these residues on the encapsidation-replication dimer conformation of the influenza C polymerase (Carrique et al. 2020) (Figure 3A-B). These paired residues are located within the N-terminal and 627 domains of PB2 and within the C-terminal domain of PA (Figure 3B, C). The residues PB2 591 and PB2 627 interact in the encapsidation-replication dimer conformation of the polymerase with host protein ANP32A (Carrique et al. 2020) (Figure 3A), and mutations in these residues are known to cooperatively increase polymerase activity in H1N1 viruses (Mehle and Doudna 2009; Liu et al. 2012). PB2-645 and PB2-199 are located near PB2-591 and PB2-627 and ANP32A and thus could cooperate with these residues to modify ANP32A binding and replication. Thus, our wMI approach identified a known cooperative interaction and at least two other interactions that are structurally plausible.

**Figure 3.**
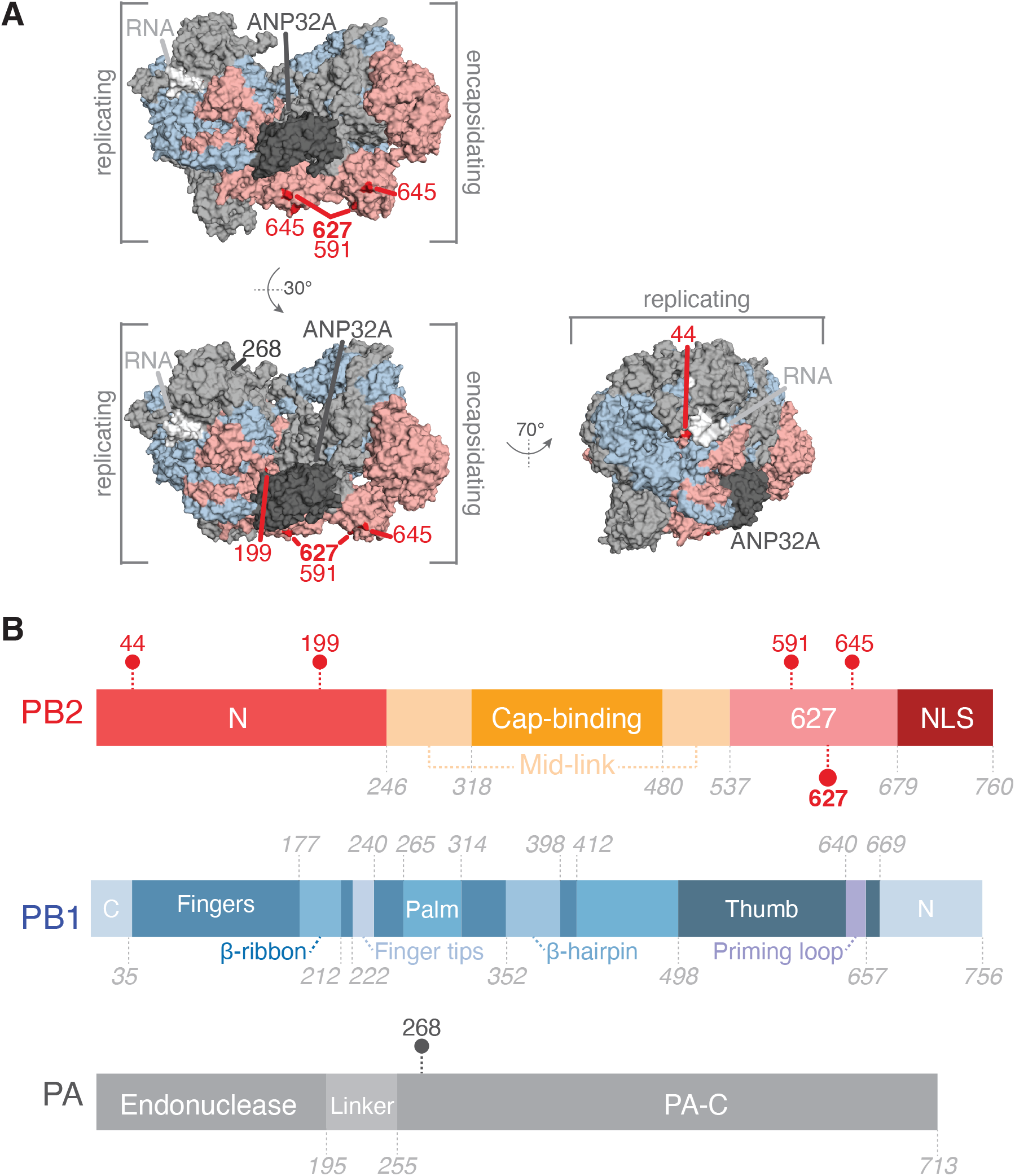
Coevolving residues with PB2-627. (A) Residues that coevolve with PB2-627 shown highlighted on the replicating-encapsidating dimer conformation of the Influenza C polymerase (PDB ID 6XZR) (Carrique et al. 2020). Highlighted residues are in dark red (PB2) or dark grey (PA). (B) Domain organization of the IAV polymerase with coevolving residues indicated.

We plotted the changes in residue frequency for PB2 627 and the five wMI-paired residues to identify the specific substitutions that account for the wMI score. These plots reveal co-incident mutations around 2011 (Figure S1) that likely underlie the wMI signal. Interestingly, one of these mutations is PB2 K627E, a reversion of the human adaptive PB2 E627K. The sequence metadata for all sequences containing this reversion revealed that the co-mutations underlying the wMI arose from a cluster of human infections in the United States Midwest with swine-derived vH3N2 viruses containing the M segment of H1N1/pdm2009. The shared PB2 mutations we identified in these viruses also suggest a possible reassortment event with PB2, which is further supported by the proximity of these residues to the binding site of host ANP32A. In all, this analysis demonstrates that wMI can identify distinct epidemiological features within viral sequence datasets spanning extensive periods or geographic areas.

We next examined whether the top wMI pairs (z-score > 4) represent interactions within or between the three polymerase subunits (Figure S2). Given that the polymerase subunits have similar substitution rates (Bhatt et al. 2011) and similar protein lengths, we would expect similar numbers of co-mutating residue pairs among each of the six gene segment pairs purely by chance. However, we observed that a large majority (869/2671 residue pairs) of top wMI pairs are specifically between PB2 and PA (a single category). Relatively few of the top wMI pairs involve PB1 at all (871/2671 residue pairs totaled across all segment pairs involving PB1). One explanation for this result is that H3N2 PB2 and PA have coevolved for a much longer period as they were inherited from the 1918 H1N1 virus, while PB1 was introduced through a reassortment event with an avian IAV in 1968 (Kawaoka et al. 1989). Another possible explanation is that PB2 and PA contain highly dynamic domains that together coordinate complex activities such as cap-snatching and dimerization (Te Velthuis and Fodor 2016). 33% of top wMI pairs include residues in the cap-binding domain of PB2 or the endonuclease domain of PA, both involved in cap-snatching, despite these domains only comprising 16% of the residues in the polymerase complex. This suggests that wMI captures coevolutionary interactions related to the enzymatic functions of the IAV polymerase.

### wMI networks identify higher order coevolutionary relationships

The wMI statistic captures coevolutionary relationships between pairs of residues. However, the coevolutionary relationships that drive polymerase function may involve more than two residues. Thus, we constructed wMI networks to extend the inherently pairwise MI statistic to encompass relationships among larger groups of residues. In these networks, nodes represent residues, and edges represent the normalized wMI (z-score) between residues.

When a network is generated with an edge for each of the top wMI pairs (n = 2671), the resulting visualization is dense and challenging to interpret due to the high degree of interconnectedness within the network. Therefore, we sought an approach to focus on the most important higher order wMI relationships within our data. Percolation theory states that in a random network, one giant interconnected graph (as opposed to many small isolated subgraphs) will quickly form as the probability of drawing an edge is increased (Newman 2018). Given that random networks tend toward a giant subgraph, we identified an edge-strength (normalized wMI) threshold at which the behavior of our network is most distinct from one containing a giant subgraph. In other words, since a network with a giant subgraph is characterized by one large subgraph with many nodes and few other subgraphs, we set our threshold to minimize the size of the largest subgraph relative to the average size of all other subgraphs (i.e. the relative maximum subgraph size, see Figure S3) (Strayer et al. 2023). This threshold results in a network visualization containing nine distinct subgraphs encompassing relationships among 40 residues (Figure 4A).

**Figure 4.**
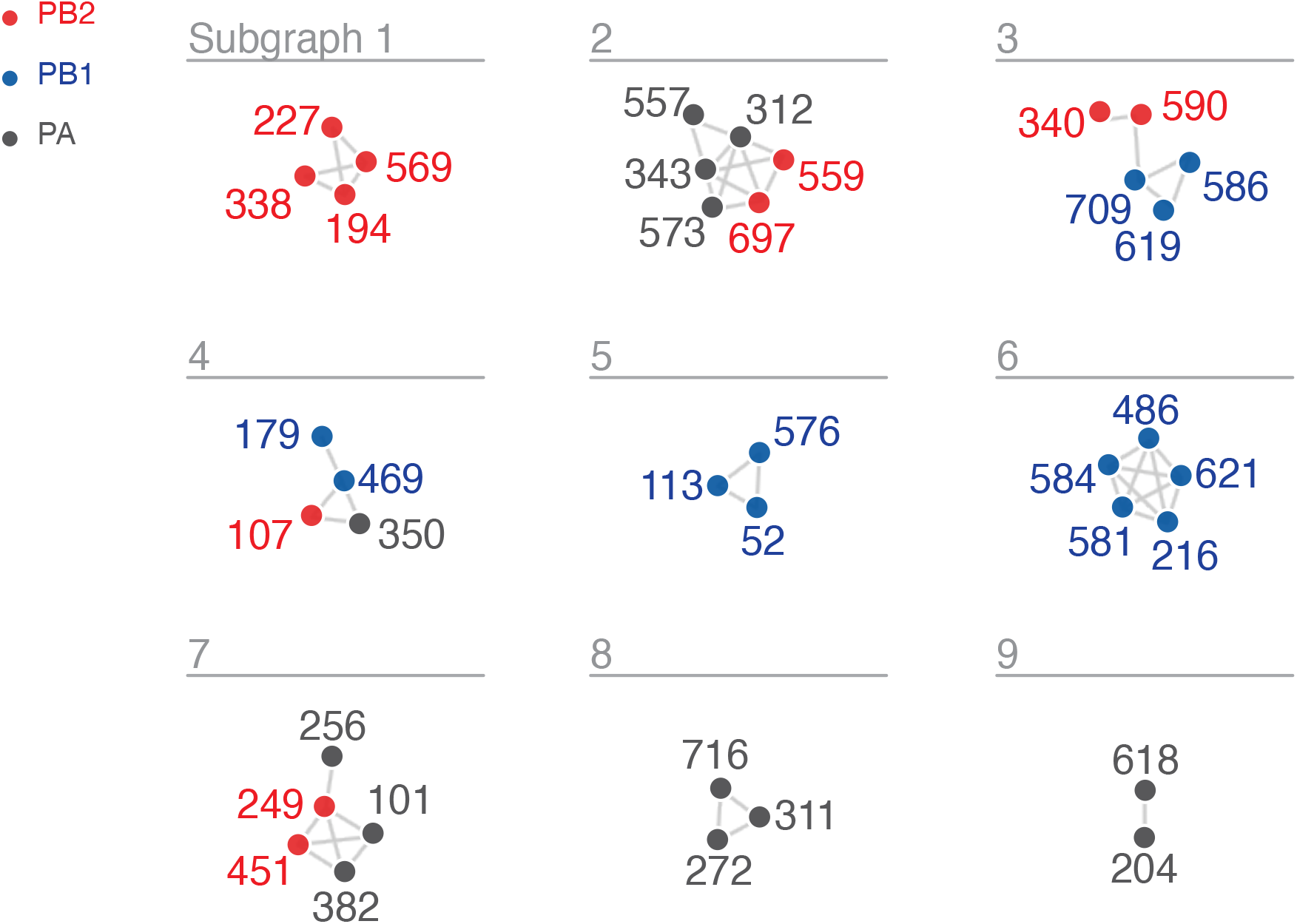
wMI network of the H3N2 polymerase (PB2, PB1, PA). Nodes represent residues and edges repre- sent the normalized wMI (z-score) between residues. Residue nodes are colored red for PB2, blue for PB1, and grey for PA. An edge threshold was set at the normalized wMI score (58) that minimizes relative maximum subgraph size (see Figure S3). The network visualization was created using the associationsubgraphs package for R (Strayer et al. 2023).

We investigated the residues within the first two subgraphs to identify potential mechanisms behind their coevolution. Subgraph 1 contains four residues within PB2: 194, 227, 338, and 569. Plotting the changes in amino acid frequency for these residues reveals that a selective sweep starting around 1985 (Q194R, M227I, I338V, and T569A) explains much of the co-evolutionary signal (Figure 5A). The location of these residues on the replication-encapsidation polymerase structure (Carrique et al. 2020) suggests that they may participate in dimerization and binding of host ANP32A; residues 194, 227, 338, and 569 are located in the dimerization interface of the RNA-bound replicating polymerase, and residue 569 is near the host ANP32A binding site (Figure 5B). The mutations Q194R and V227I were also shown to be human-adaptive markers in a study of H3N2 sequences from human and avian hosts (Wen et al. 2018). Subgraph 2 contains a mix of PA and PB2 residues: PA-312, PA-343, PA-557, PA-573, PB2-559, and PB2- 697. A selective sweep around 2005 (PA R312K, PA A343S, PA M557I, PA I573V, PB2 T559A, and PB2 L697I) contributed to the high wMI among these residues (Figure 5A). These residues are located within the C-terminal domain of PA and the 627 and NLS domains of PB2, at the interface of the replication-encapsidation polymerase dimer (Carrique et al. 2020) (Figure 5C). In addition, the mutations PB2 T569A (Subgraph 1) and PB2 T559A (Subgraph 2) are known regulators of host-range expansion in the H7N9 polymerase (Chen et al. 2016). In all, the construction of wMI networks in the H3N2 polymerase identified relationships between residues that regulate host adaptibility and are likely involved in replication, encapsidation, and association with host ANP32A.

**Figure 5.**
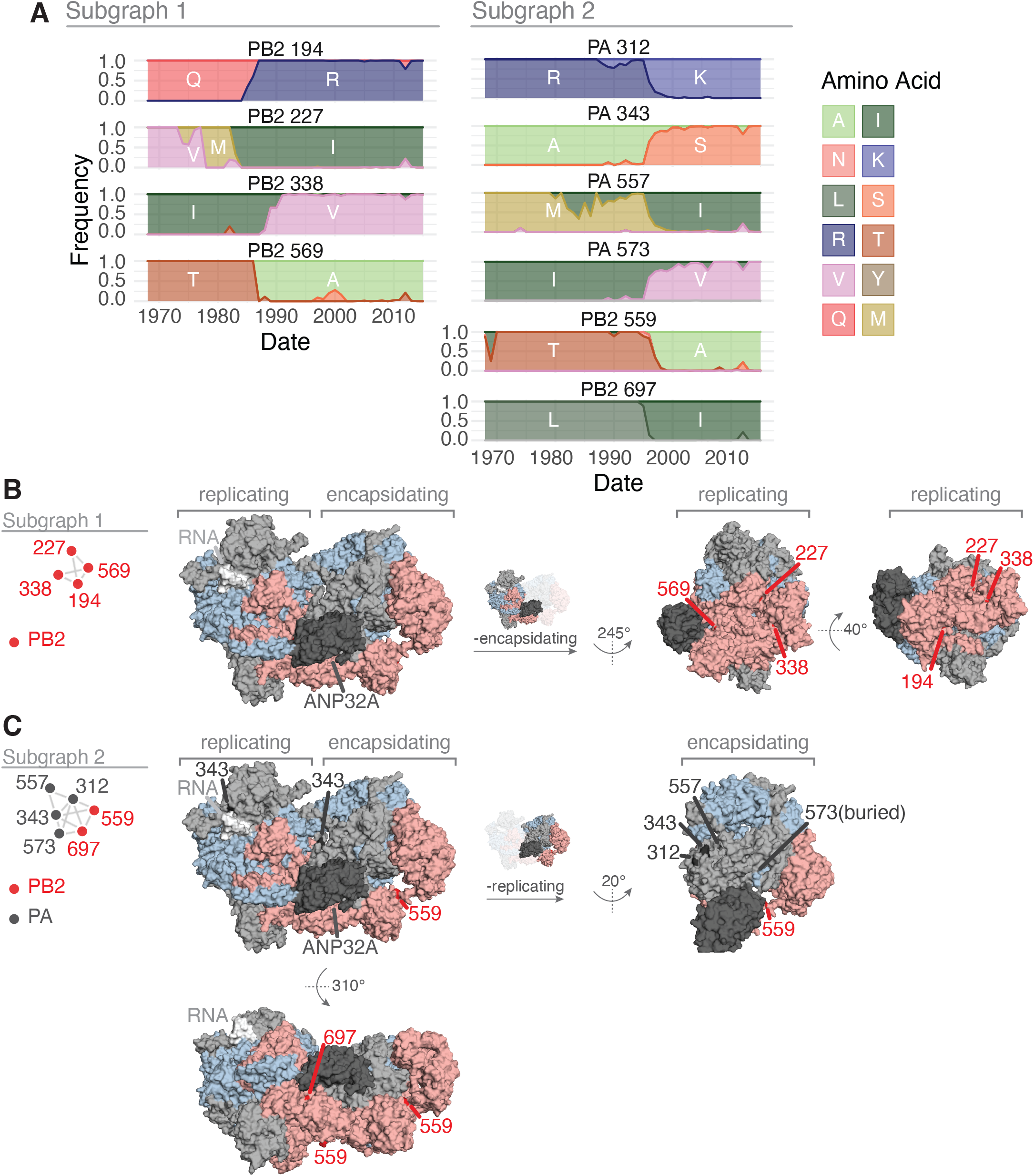
Features of the residues in Subgraphs 1 and 2 from the wMI network of the H3N2 polymerase. (A) Amino acid frequencies from 1968 – 2015 for the residues within Subgraph 1 (left) or Subgraph 2 (right). (B, C) location of the residues in Subgraph 1 (B) and Subgraph 2 (C) plotted on the replicating-encapsidating dimer conformation of the Influenza C polymerase (PDB ID: 6XZR) (Carrique et al. 2020). In B-C, highlighted residues are in dark red (PB2) or dark grey (PA).

### wMI networks can reveal genetic hitchhiking

Co-evolving sites within the IAV polymerase may be falsely assumed to have biological significance due to genetic hitchhiking with HA or NA during antigenic drift. Antigenic variants that promote immune escape are under strong selection; when these mutations undergo a selective sweep, neutral or even deleterious mutations in other regions of the IAV genome may also rise in frequency in the population due to linkage disequilibrium (Chen and Holmes 2010; Lyons and Lauring 2018). We accounted for this possibility with HA by calculating wMI scores for a joint MSA of the three polymerase proteins and HA. We found that most of the top wMI pairs (z-score > 4) occur within HA, which is expected due to the higher substitution rate of HA versus the polymerase proteins (Figure S4A) (Bhatt et al. 2011). In addition, the top wMI pairs within HA antigenic regions A-E (Wiley et al. 1981; Wilson et al. 1981; Skehel et al. 1984) have higher normalized wMI overall than top wMI HA pairs in other regions of the protein (Figure S4C). Interestingly, there are fewer top wMI pairs between the polymerase proteins than between each polymerase protein and HA (Figure S4A, B). Overall, this suggests a high level of coevolution between the polymerase complex and HA and underscores the need to parse coevolution due to functional relationships versus genetic hitchhiking due to antigenic selection.

We then constructed a wMI network and reasoned that subgraphs containing both polymerase and HA residues represent potential genetic hitchhiking events (Figure 6A and S5). In the polymerase-HA wMI network many of the relationships with HA involve residues within the antigenic regions A-E (Wiley et al. 1981; Wilson et al. 1981; Skehel et al. 1984), including known epistatic residues within antigenic region B (Figure 6) (Wu et al. 2020). As the wMI relationships between the polymerase and HA antigenic residues may indicate genetic hitchhiking, we defined a set of polymerase-only subgraphs likely to be functionally important. We again evaluated the functional implications of the residues in these networks by examining changes in amino acid frequency and placing them on the post-cap-snatching polymerase structure (Fan et al. 2019) (Figure 7A-D). Subgraphs 4 and 8 contain residues co-varying in amino acid frequency between 1970 and 2005 (Figure 7A). Subgraph 4 is a pairwise interaction between PB1-619 and PB1-709, which are located in the thumb and C-terminal domains, respectively (Figure 7B). The thumb domain forms the right-side wall of the viral RNA-dependent polymerase (RdRp) active site chamber, while the C-terminal domain interacts closely with the PB2 N-terminus and PA endonuclease domains. In addition, the mutations V709I and D619N in PB1 each lead to increased polymerase activity (by minigenome assay in human cells) in the early pandemic H3N2 strain A/Hong Kong/1/1968(HK/68) (Sun et al. 2022: 1). PB1-52 and PB1-576 of Subgraph 8 are in the finger and thumb domains of PB1 (Figure 7C). The finger domain of PB1 forms the roof and left-side wall of the RdRp active site chamber. While PB1-52 and PB1-576 are in not in close proximity, the mutation PB1 I576L is one of seven differences between consensus avian PB1 and H1N1 PB1 from the 1918 pandemic (Taubenberger et al. 2005), and K52R is found in a significantly higher proportion of IAVs isolated from humans than swine (Chen et al. 2017). Thus, PB1-52 and PB1-576 may be residues associated with host adaptability. Subgraph 10 contains residues from all three polymerase subunits: PB2-107, PB1-469, and PA-350. These residues undergo two collective shifts in amino acid frequency, first starting in 1977 and again near 1996 (Figure 7A). They are located in the N-terminal domain of PB2, the palm domain of PB1, and the C-terminal domain of PA (Figure 7D). The N-terminal domain of PB2 closely associates with the RdRp, and the C-terminal domain of PA associates with the thumb domain of the RdRp. The PB1 palm subdomain forms the floor of the RdRp active site chamber. Residue PB1 469 is also a determinant of host range for H1N1: the mutation A469T determines transmissibility in guinea pigs, and this mutation also arose after serial passage of pdm09 H1N1 in pigs (Wei et al. 2014). In all, the non-HA-associated subgraphs highlight residues near the main enzymatic activities of the RdRp that may alter host adaptability.

**Figure 6.**
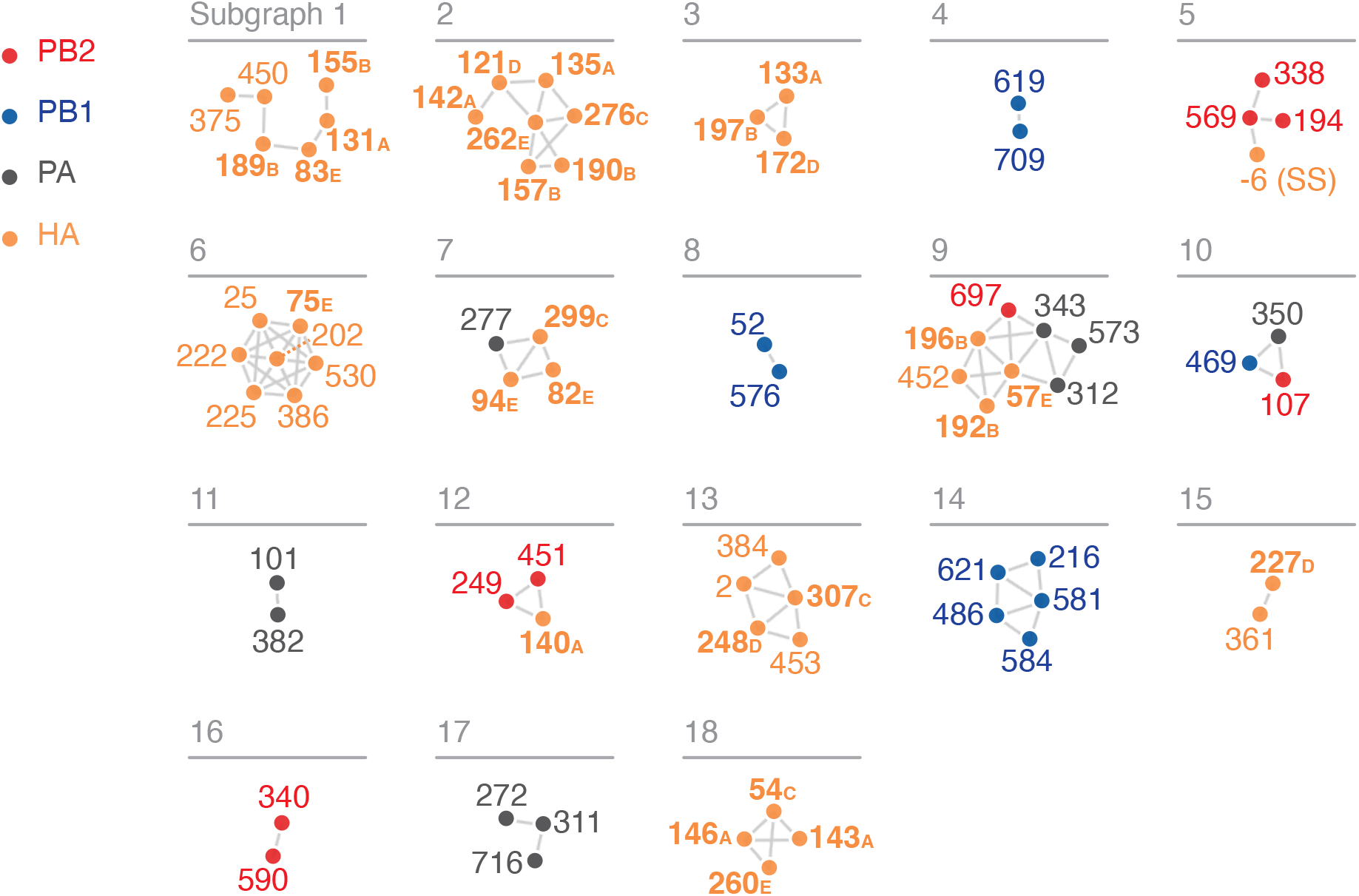
wMI network of the H3N2 polymerase (PB2, PB1, PA) and HA. Nodes represent residues and edges represent the normalized wMI (z-score) between residues. Residue nodes are colored as in Figure 4, plus orange for HA. HA residues that are located in antigenic regions A-E are shown in **bold**. Residue -6 (SS) is in the cleaved N-terminal signal sequence of HA. An edge threshold was set at the normalized wMI score (40.506) that minimizes relative maximum subgraph size (see Figure S5). The network visualization was created using the associationsubgraphs package for R (Strayer et al. 2023).

**Figure 7.**
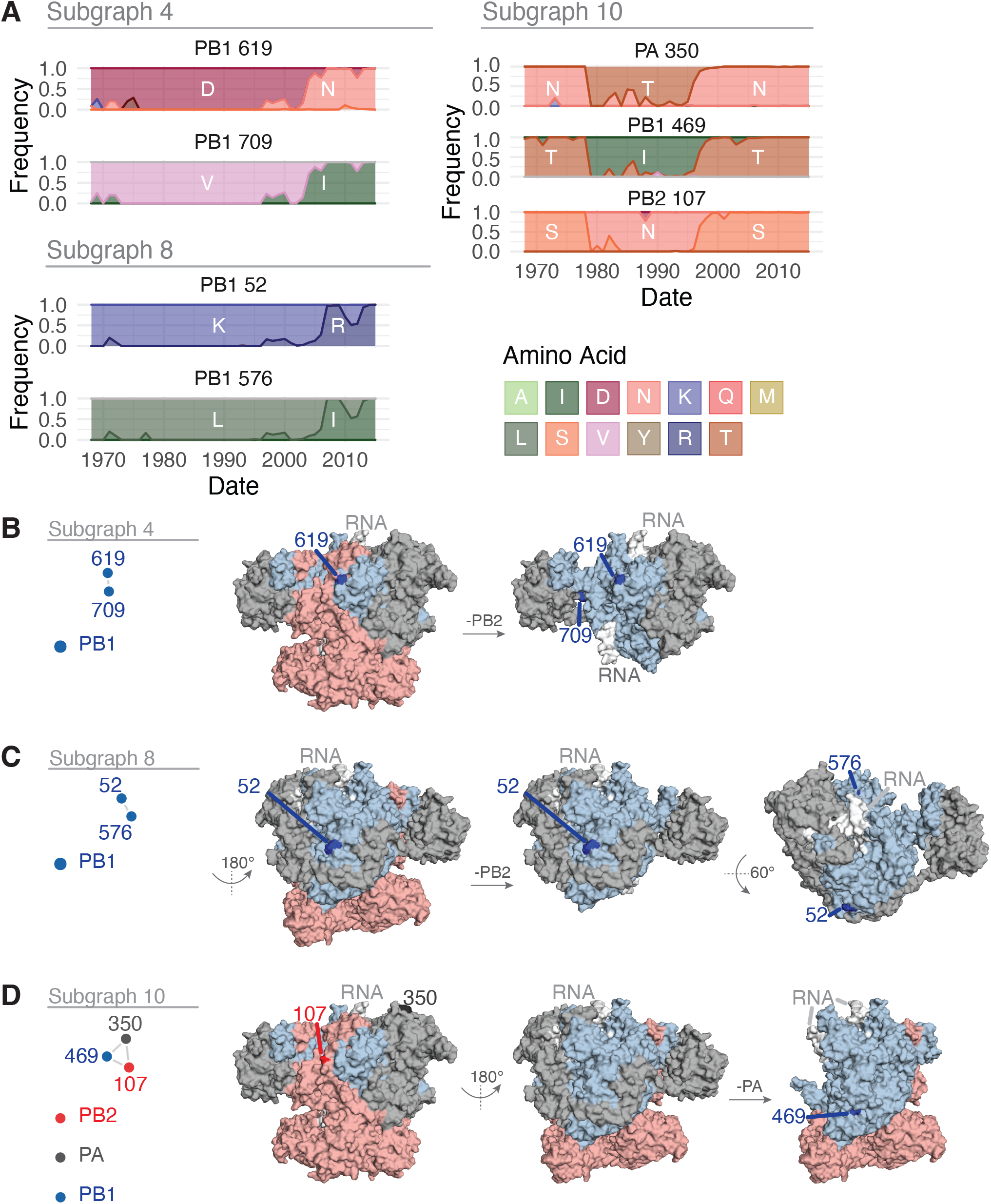
Features of the residues in Subgraphs 4, 8, and 10 from the wMI network of the H3N2 polymerase and HA. (A) Amino acid frequencies from 1968 – 2015 for the residues within Subgraph 4 (top left), Subgraph 8 (bottom left), or Subgraph 10 (right). (B-D) location of the residues in Subgraph 4 (B), Subgraph 8 (C), or Subgraph 10 (D) plotted on the post-cap-snatching conformation of the H3N2 polymerase (PDB ID: 6RR7) (Fan et al. 2019). In B-D, highlighted residues are in dark red (PB2), dark blue (PB1), or dark grey (PA).

Subgraphs that contain both polymerase and HA residues represent potential genetic hitchhiking. However, the presence and direction of hitchhiking must be investigated case-by-case and confirmed experimentally. For example, polymerase residues in subgraphs 5 and 9 from the polymerase-HA network (corresponding to subgraphs 1 and 2 in the polymerase-only network) may have high wMI due to genetic hitchhiking with mutations in HA. The residues in subgraph 5 all underwent a selective sweep around 1985 (Figure S6). However, the mutation in PB2-194 precedes the mutation in HA (-6). Thus, whether genetic hitchhiking is occurring, and the direction of potential hitchhiking, is unclear. On the other hand, the residues in subgraph 9 underwent a simultaneous selective sweep starting in 1995. The timing of this sweep and the association with residues in HA antigenic regions B and E indicate that high wMI among polymerase residues in this subgraph may be due to selection acting on mutations in HA. In all, wMI networks are a useful diagnostic tool to form hypotheses about hitchhiking relationships that may be further investigated.

## Discussion

The wMI metric introduced in this study addresses several issues using information-based measures to investigate evolution and coevolution in rapidly evolving populations. Weighting across years accounts for sampling variations over time. Using the wMI metric, we identified a robust coevolutionary relationship between PB2-627 and PB2-591. These residues are known to interact and are essential for host range expansion (Mehle and Doudna 2009), validating our approach. We generated network visualizations (Newman 2018; Strayer et al. 2023) of wMI to facilitate the identification of higher-order interactions and provide a method for addressing genetic hitchhiking. This analysis identified clusters of coevolving residues with roles in cap- snatching, dimerization, replication, and host adaptability. We included HA in the network to identify polymerase-only wMI relationships with potential roles in the enzymatic functions of the RdRp and host-range expansion.

The wMI method has several strengths compared to other methods for detecting coevolution. Unlike weighting by sequence similarly, wMI preserves the changes in allele frequency that are crucial for detecting coevolution in rapidly evolving populations. wMI also does not require fitting a model and thus does not suffer from model selection or fit issues (Dutheil 2012). In addition, the simplicity of the wMI metric makes it relatively easy and fast to implement. Previous methods to detect coevolution have used intra-molecular distances from structural data as a benchmark (Weigt et al. 2009; Morcos et al. 2011; Kamisetty et al. 2013; Figliuzzi et al. 2016). However, structural proximity is only one factor that can lead to coevolution (Ackerman et al. 2012). Other factors include protein function, RNA function, RNA structure, stochastic processes, and phylogeny (though corrected in our approach). In one recent study, structural proximity was found to contribute to general low-level MI across a protein, while functional relationships are indicated by strong MI (Mohan et al. 2022). Thus, evaluating coevolution- detection methods based on structure alone will select methods that cannot capture all of the biology at play.

The wMI method combined with network visualization provides a new way of identifying and excluding co-evolutionary relationships due to genetic hitchhiking. Our method cannot definitively confirm or refute the presence of hitchhiking. However, it does produce a tractable set of hypotheses about whether hitchhiking underlies the most important coevolutionary relationships in a system. Accounting for other proteins besides HA under strong selection, such as NA, will yield additional insights into hitchhiking relationships between IAV genes.

The methods introduced in this study have several limitations. A primary limitation is that increasing (through weighting) the influence of a year with few observations can increase the variability in the resulting wMI. However, assuming there is no pattern in sampling variability, the sum effect on the wMI of up-weighting all low-observation years should be negligible. A second limitation of using wMI is the assumption that there are no unknown confounders. We assume in this method that the sequence observations in each year represent a random sample of the viral genomes present in that year. In recent decades, the distribution of sequences from different geographic regions has become heavily biased towards North America and Europe. However, increased spread of IAVs between geographic regions (Grais et al. 2003) means the effect of this bias on genome variability is reduced. Furthermore, the wMI method introduced in this paper could be similarly used to address uneven sampling across geographic regions, though the assumptions inherent in equal weighting (versus incidence-weighting) may prove problematic. An additional possibility is that even if the genomes in our dataset represent a sufficiently random sample, the observed changes in allele frequency could reflect genetic drift rather than natural selection. Another major limitation of this study is that MI and wMI can only detect dependencies among residues that have evolved, as residues that are fully conserved over the study period will have an entropy of 0. Thus, MI and wMI cannot capture the assuredly meaningful relationships among strongly conserved positions. A final limitation is that wMI does not provide insight into the coevolving residues’ function(s). Instead, relationships identified using wMI represent preliminary hypotheses for further investigation.

The wMI-edge threshold we set is informed by the behavior of random networks and helps form hypotheses about functional coevolutionary relationships to test experimentally (Newman 2018; Strayer et al. 2023). However, coevolutionary relationships within the H3N2 polymerase are not limited to what happens at that threshold. Furthermore, it is not easy to interpret how a particular threshold influences the findings. For example, would a slightly higher or lower threshold result in different hypotheses regarding hitchhiking? To this end, we have developed a Shiny Application (Chang et al. 2022) to dynamically visualize our network at different thresholds. The Shiny Application can be accessed at https://virusevolution.shinyapps.io/MI_Networks_App/ and contains all the wMI results presented in this study.

The wMI metric we introduce solves several critical issues that have limited the application information-theoretic methods to studies of evolution. The simplicity of the wMI metric means that implementation and application to other systems is relatively straightforward. Notably, the solutions we propose for uneven sampling, identifying higher-order interactions, and accounting for genetic hitchhiking, have utility in systems beyond the H3N2 polymerase.

## Materials and Methods

### H3N2 Sequence Acquisition

IAV polymerase sequences (amino acid) were downloaded from GISAID on August 30^th^, 2022. Entries were filtered for A/H3N2 subtype, human host, all locations and collection times, and only complete sequences for PB2, PB1, PA, and HA. In all, this resulted in 7250 entries. Each segment was downloaded as a separate fasta file. Metadata for each sequence is provided in Supplemental Table 1 (GISAID acknowledgement table).

### H3N2 Sequence processing

Sequences were first filtered to remove any entries passaged in egg, using the regular expression “egg|Egg|E[0-9]|AM|Am|E”. Entries with duplicate sequences were removed entirely. Lastly, any sequences containing insertions or deletions were removed by filtering for sequence length. In all, these filtering steps removed 314 sequences, leaving 6936 sequences for downstream analysis. Sequences for all segments were then aligned using MAFFT v7.490 (Katoh et al. 2002) (released October 30^th^, 2021). Aligned sequences were concatenated by isolate ID using the order: PB2-PB1-PA-HA.

### Calculation of weighted amino acid frequencies, entropy, and mutual information

Weighted amino acid frequencies were calculated according to Equations 3 – 5. Shannon entropy and mutual information were calculated according to the Equations 1 and 2. For weighted entropy and mutual information, the weighted amino acid frequencies were used in Equations 1 and 2 as described above. Finally, each MI or wMI was corrected for the influence of phylogenetic signal using the average product correction (Equation 6) as previously described(Dunn et al. 2008).

### Sliding-window analysis

5-year sliding windows starting at each year were constructed from 1968 – 2011, moving over one year per window (the last window including years 2011 – 2015). MIs (including the average product correction) were calculated for the sequences in each window using Equations 1 and 2.

### SARS-CoV-2 Spike simulations

Washtenaw County SARS-CoV-2 sequences were downloaded (as nucleotide) from GISAID using the isolate IDs provided in Supplemental file 2 (GISAID acknowledgement table).

Sequences were subset to the spike gene and filtered to remove sequences containing “N” or “K”. Then the sequences were aligned using MAFFT v7.490 (Katoh et al. 2002) (released October 30^th^, 2021), and aligned sequences were translated into amino acids and subset to the RBD domain. Spike RBD sequences were then sampled by month with replacement using the binned sampling density of the IAV polymerase (number of bins = 12, to match the number of months in the Spike RBD dataset). This sampling was repeated to generate 100 independent samples. The raw MI, weighted MI (using disease incidence), and weighted MI (using 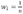) were calculated for each sample, and the raw MI was calculated for the original Spike RBD dataset. The original sampling frequency is roughly proportional to disease incidence (Figure 2A). The average product correction was not applied to any of the calculated MIs in this analysis. The Spearman correlation was then calculated to compare the original Spike RBD dataset MI to the MIs for each sample. A kernel density plot was generated using the geom_density function with default parameters from the ggplot2 R package (Wickham 2016). Washtenaw County SARS-CoV-2 positive cases were taken from “Cases and Deaths by County by Date of Onset and Date of Death” downloaded from https://www.michigan.gov/coronavirus/stats

### Network construction and thresholding

Networks were visualized using the associationsubgraphs package for R(Strayer et al. 2023). The input data was subset to the top (length of MSA / 2) pairs to improve computational speed and rendering. A network edge threshold was chosen using the “min-max rule” (ie, minimizing the relative maximum subgraph size) as previously described (Strayer et al. 2023).

### Protein Visualizations

All protein visualizations were constructed using PyMOL (version 2.5.4)(Anon). Python scripts to generate PyMOL image files were adapted from scripts on the Bloom lab github site (https://github.com/jbloomlab/PB2-DMS) (Soh et al. 2019). Domain structure for the polymerase proteins was adapted from (Pflug et al. 2014).

### Code Availability

Code for generating all analyses and visualizations in this study is provided at https://github.com/lauringlab/timeMI. Functions for calculating raw and weighted MI are available as a separate package for R: https://github.com/lauringlab/weightedMI. Interactive network visualizations containing all wMI results published in this study can be viewed at https://virusevolution.shinyapps.io/MI_Networks_App/

## Acknowledgements

We thank Dirk Eggink for helpful discussion. This work was supported by NIH R01 AI170520 (to ASL) and a Burroughs Wellcome Fund Investigator in the Pathogenesis of Infectious Disease Award (to ASL). Dr. Arcos was supported, in part, by NIH T32 AI007528.

