## Supplemental Figures for "Mutual information networks reveal evolutionary relationships within the influenza A virus polymerase"

**Figure S1**

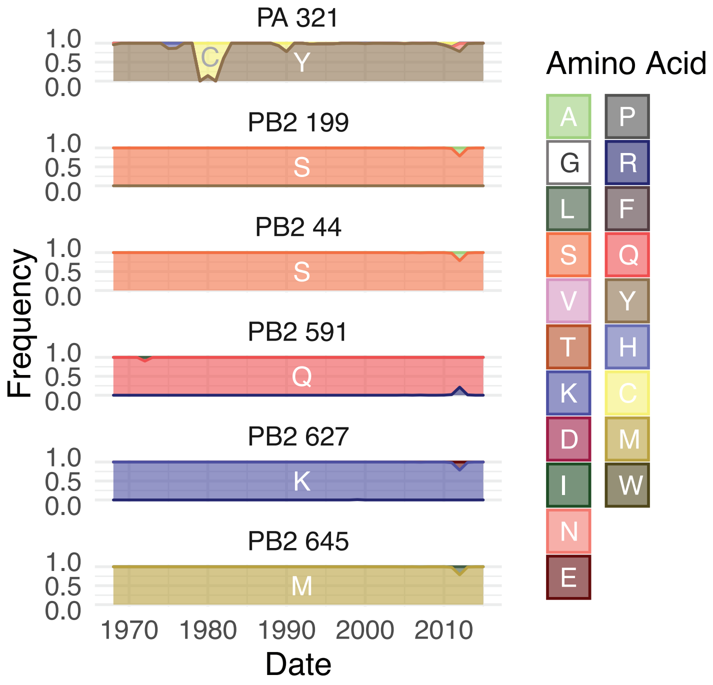


**Figure S1.** Amino acid frequencies from 1968 – 2015 for the top wMI residue pairs with PB2-627, related to Figure 3.

**Figure S2**

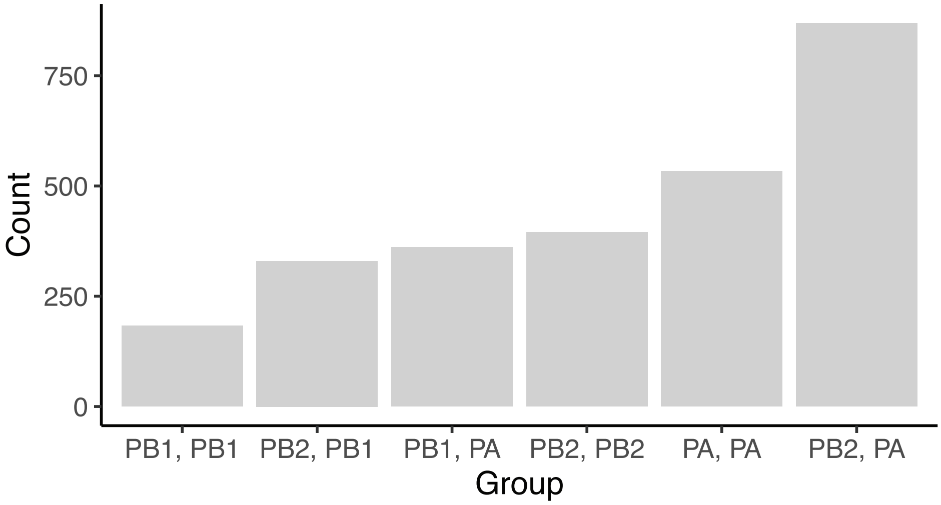


**Figure S2.** The number of top (z-score > 4) wMI residue pairs within and between the three subunits of the H3N2 polymerase (PB2, PB1, PA). The x-axis is arranged by count. Related to Figures 3 – 5.

### Figure S3

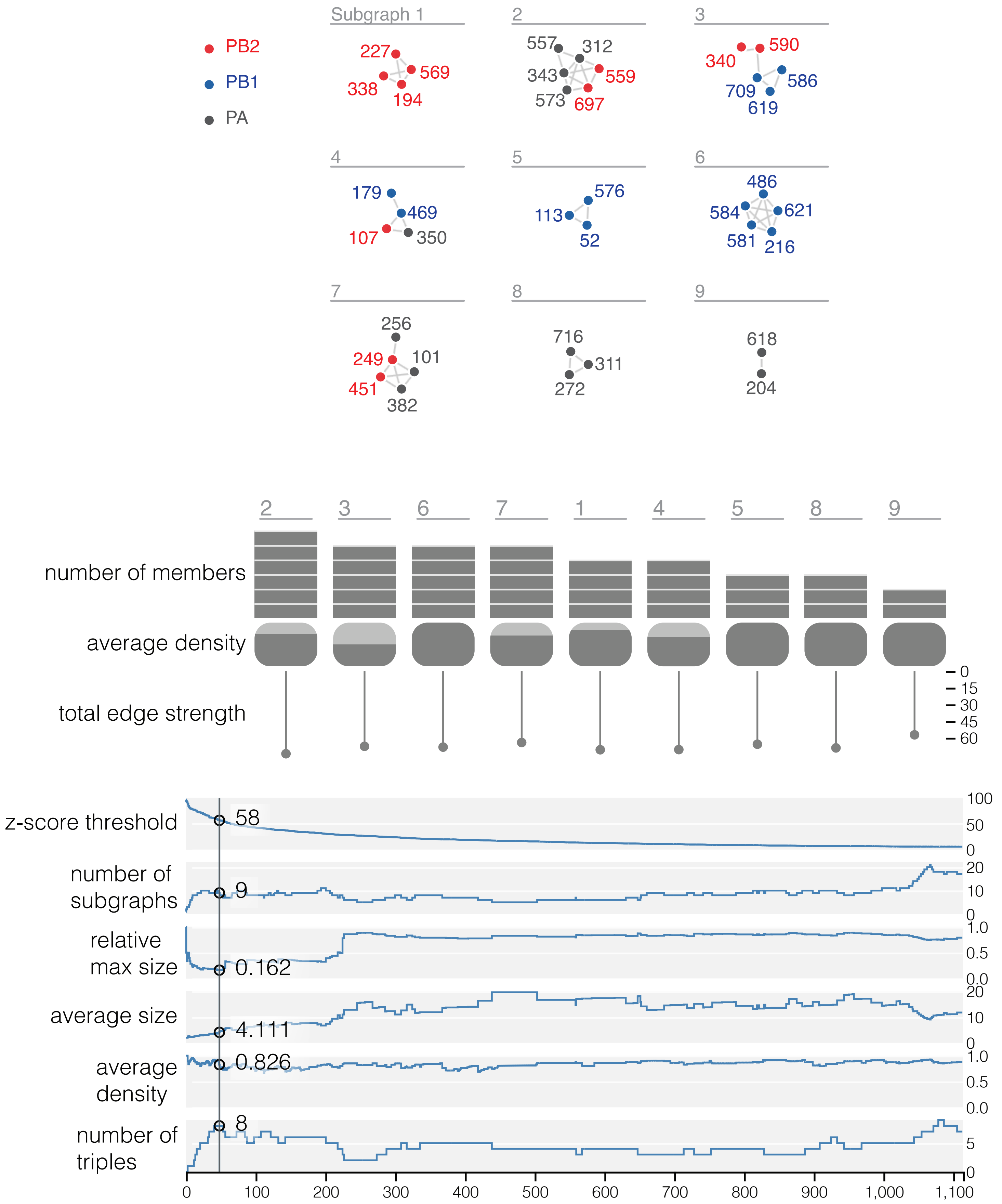

PB2
PB1
PA
Subgraph 1
227
569
338
194
2
557
312
343
559
573
697
3
340
590
586
709
619
4
179
469
107
350
5
576
113
52
6
486
584
621
581
216
7
256
249
101
451
382
8
716
311
272
9
618
204
2 3 6 7 1 4 5 8 9
number of members
average density
total edge strength
0
15
30
45
60
z-score threshold
58
100
50
0
number of
subgraphs
9
20
10
0
relative
max size
0.162
1.0
0.5
0.0
average size
4.111
20
10
0
average
density
0.826
1.0
0.5
0.0
number of
triples
8
5
0
0 100 200 300 400 500 600 700 800 900 1,000 1,100

**Figure S3.** Network diagnostics for the wMI network of the H3N2 polymerase. Top, network visualization of the wMI scores within the H3N2 polymerase. Nodes represent residues and edges represent the normalized wMI (z-score) between residues. An edge threshold was set at the normalized wMI score (z-score >= 58) that minimizes relative maximum subgraph size. Middle, plots showing the number of members, average density, and total edge strength of the indicated subgraphs. Density refers to the number of edges in a subgraph divided by the total number of possible edges in that subgraph (possible edges = (n*(n-1))/2,where n = number of nodes). Bottom, diagnostic traces of network parameters as the z-score threshold (top trace) is decreased: number of subgraphs, relative max size, average size, average density, and number of triples. Relative max size is calculated by dividing the size (number of nodes) of the largest subgraph by the number of nodes in the remaining subgraphs at a given threshold. The threshold which minimizes the relative max subgraph size is indicated by the vertical line. The network visualization and diagnostic traces were created using the associationsubgraphs package for R (Strayer et al. 2023). Related to Figure 4.

**Figure S4**

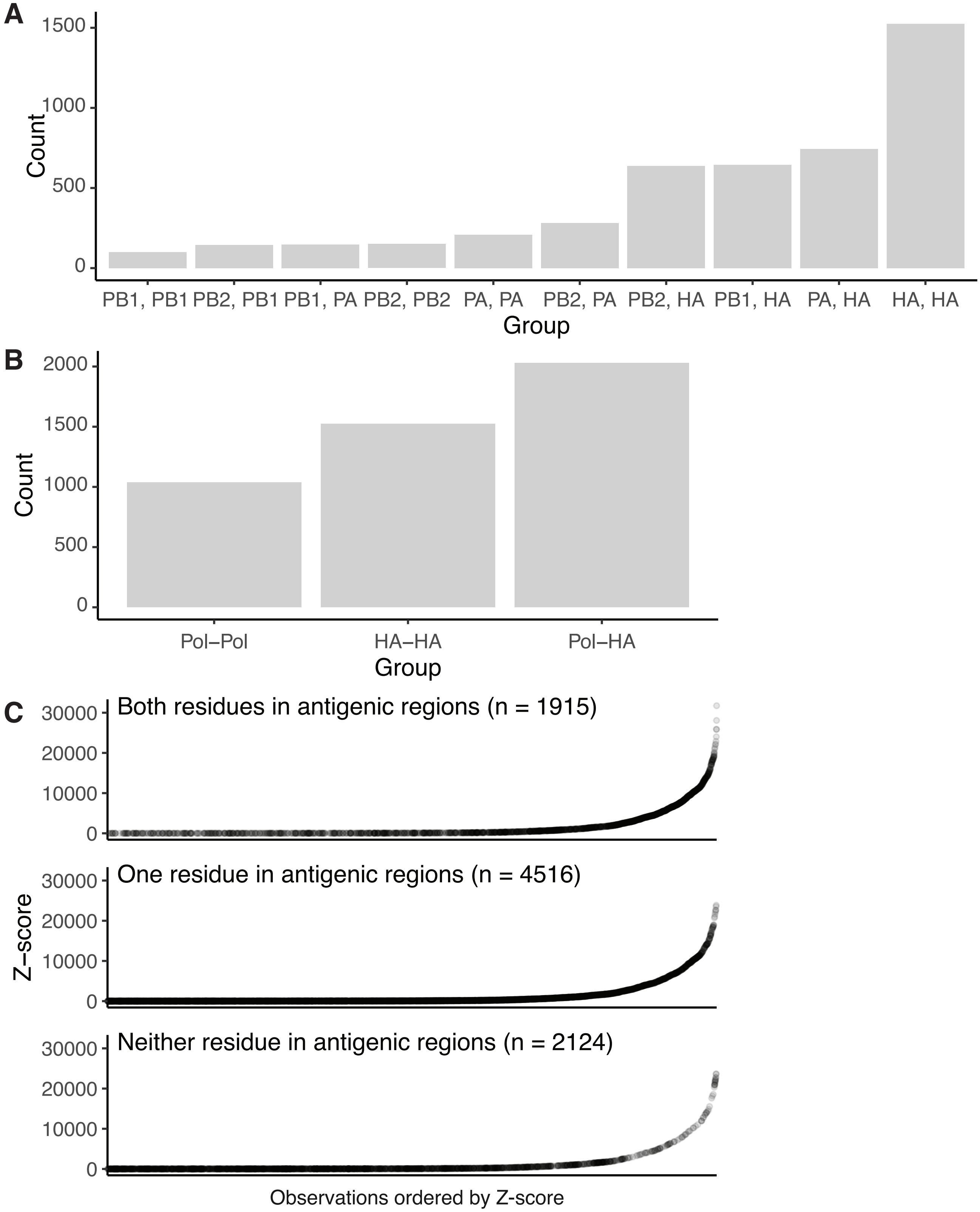


**Figure S4.** wMI residue pairs within and between the H3N2 polymerase and HA. (A) Bar chart showing the number of top (z-score > 4) wMI residue pairs within and between each H3N2 polymerase subunit (PB2, PB1, PA) and HA. The x-axis is arranged by count. (B) Summarized version of the bar chart in (A), where the polymerase subunits are considered together as a group. (C) Plots of HA-only pairs by z-score, where top HA-only wMI pairs (z-score > 4) are grouped based on whether both, one, or neither of the residues in the pair are located in HA antigenic regions A-E. Related to Figures 6 and 7.

**Figure S5**

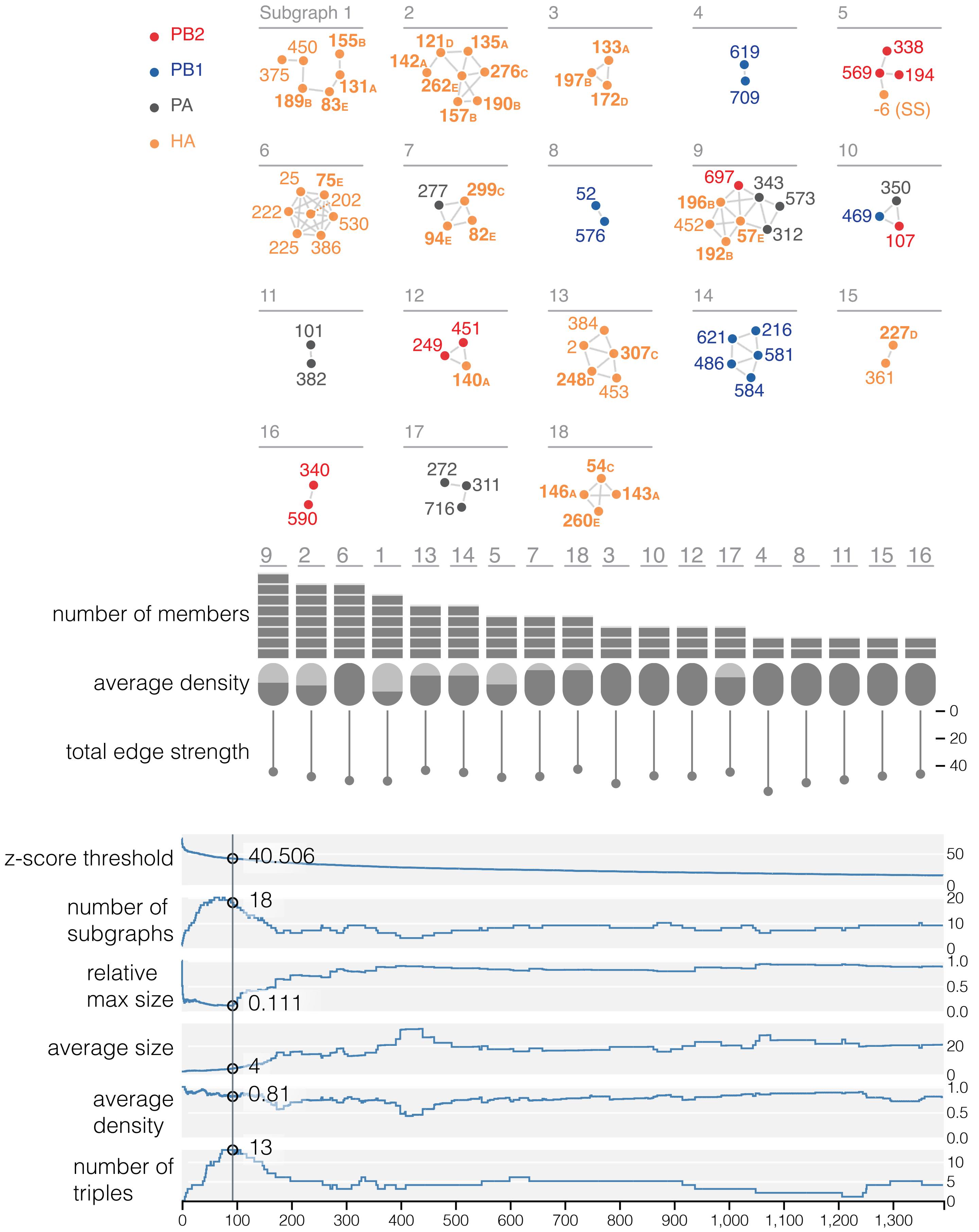

PB2
PB1
PA
HA
Subgraph 1
450
155B
375
131A
189B
83E
2
121D
135A
142A
262E
276C
157B
190B
3
133A
197B
172D
4
619
709
5
338
569
194
-6 (SS)
6
25
75E
222
202
530
225
386
7
277
299C
94E
82E
8
52
576
9
697
343
573
196B
452
57E
312
192B
10
350
469
107
11
101
382
12
451
249
140A
13
384
2
307C
248D
453
14
621
216
486
581
584
15
227D
361
16
340
590
17
272
311
716
18
54C
146A
143A
260E
9 2 6 1 13 14 5 7 18 3 10 12 17 4 8 11 15 16
number of members
average density
total edge strength
0
20
40
z-score threshold
40.506
50
0
number of subgraphs
18
20
10
0
relative max size
0.111
1.0
0.5
0.0
average size
4
20
0
average density
0.81
1.0
0.5
0.0
number of triples
13
10
5
0
0
100
200
300
400
500
600
700
800
900
1,000
1,100
1,200
1,300

**Figure S5.** Network diagnostics for the wMI network of the H3N2 polymerase and HA. Top, network visualization of the wMI scores within the H3N2 polymerase. Nodes represent residues and edges represent the normalized wMI (z-score) between residues. Residue -6 (SS) is in the N-terminal cleaved signal sequence of HA. HA residues that are located in antigenic regions A-E are shown in **bold**. An edge threshold was set at the normalized wMI score (z-score >= 40.506) that minimizes relative maximum subgraph size. Middle, plots showing the number of members, average density, and total edge strength of the indicated Subgraphs. Density refers to the number of edges in a subgraph divided by the total number of possible edges in that subgraph (possible edges = (n*(n-1))/2,where n = number of nodes). Bottom, diagnostic traces of network parameters as the z-score threshold (top trace) is decreased: number of subgraphs, relative max size, average size, average density, and number of triples. Relative max size is calculated by dividing the size (number of nodes) of the largest subgraph by the number of nodes in the remaining subgraphs at a given threshold. The threshold which minimizes the relative max subgraph size is indicated by the vertical line. The network visualization and diagnostic traces were created using the associationsubgraphs package for R (Strayer et al. 2023). Related to Figure 6.

### Figure S6

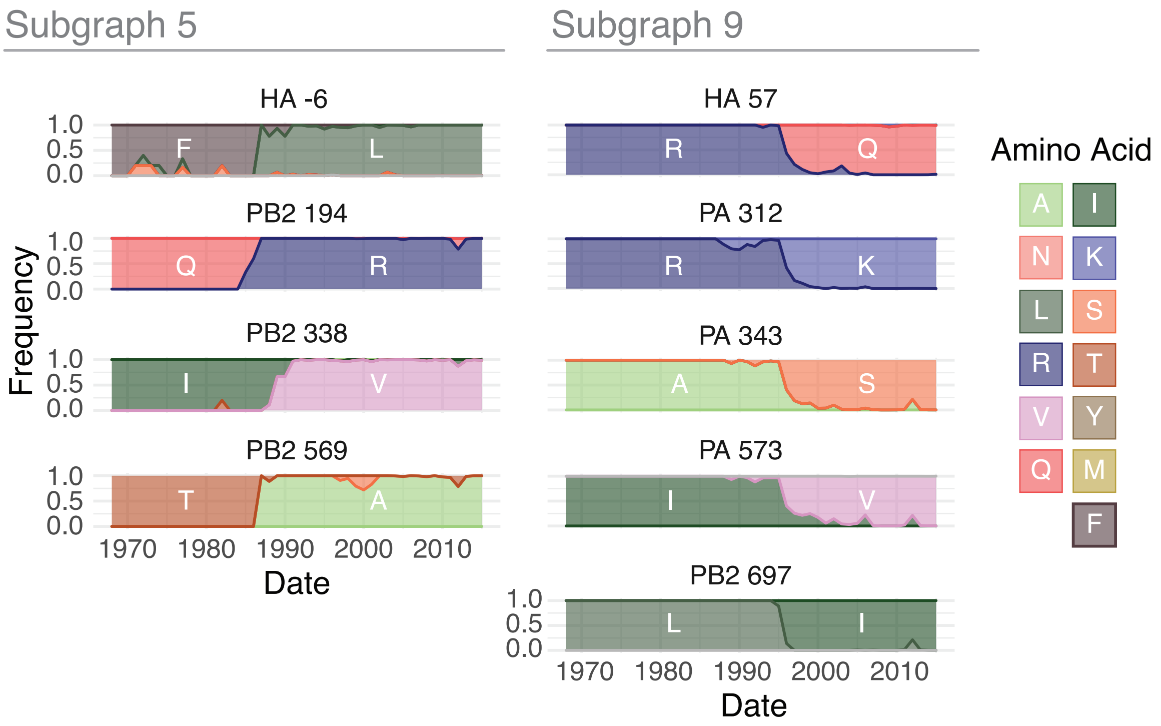


**Figure S6.** Amino acid frequencies from 1968 – 2015 for the wMI residues in Subgraph 5 (left) and 9 (right). For simplicity, a representative HA residue is shown from Subgraph 9. Related to Figure 6.
